# How to prepare the input data and run MCScanX efficiently?

**DOI:** 10.1101/2025.07.29.666888

**Authors:** Xi Zhang, David Roy Smith

## Abstract

The protocol by Wang et al. is particularly useful as it outlines the steps for efficiently identifying colinear blocks in intra-and inter-species BLASTP outputs by using MCScanX. We recently discovered that the protocol lacks the pre-processing steps for checking if there are multiple isoforms derived from alternative splicing. Conserved sequences derived from alternative splicing can have similar functional domains, to avoid mis-prediction of gene duplicates, especially for the genome data from NCBI or other online resources. Without this step, the number of duplicate genes will be overrepresented. Besides we shared some useful experience to faster preparing the input data and easier running MCScanX. This is including alternative options to prepare the ‘.gff’ input file and iterated all-against-all BLASTP processing. Lastly, we want to raise awareness of the potential challenges when preparing the input files and highlight potential issues when using the protocol.

## Main Text

The protocol provided by Wang et al. (Wang et al., 2024) is particularly useful as it outlines the steps for efficiently identifying colinear blocks in intra- and inter-species BLASTP outputs using MCScanX (Wang et al., 2012). They investigated data downloaded directly from NCBI and demonstrated the usefulness of their protocol for finding and visualizing colinear blocks in evolutionary analyses. Their downstream findings revealed MCScanX is not only acted as a stand-alone method for detecting colinear blocks, but it also can be incorporated into third-party tools to make further use of the MCScanX results. For example, SynVisio (Bandi and Gutwin, 2020) can explore the results of MCScanX and visualize them into different plots; another popular tool called GENESPACE (Lovell et al., 2022) can utilize MCScanX as the dependency for building a synteny-based pan-genome analysis; and the DupGen_finder (Qiao et al., 2019) can identify and classify different modes of duplicated gene pairs based on the built-in MCScanX algorithm.

Part 2 of the protocol by Wang et al. includes steps for downloading data directly from NCBI and preparing the necessary ‘.gff’ and ‘.blast’ files yielded from blast all-vs-all. We recently discovered that the Part 2 lacks the pre-processing step for checking if there are multiple isoforms derived from alternative splicing. Indeed, when analyzing data downloaded directly from NCBI, it can be crucial to filter out the longest transcript as the primary protein sequence.

Conserved sequences derived from alternative splicing can have similar functional domains. To avoid misprediction of gene duplicates, especially for genome data from NCBI or other online resources, it can be helpful to include a primary protein filtering step (Zhang et al., 2023, Zhang et al., 2025). This is particularly true when analyzing datasets of species with large numbers of duplicate genes. For example, doubletrouble (Almeida-Silva and Van de Peer, 2025), which is an R/Bioconductor package for gene and genome duplications analysis, uses AAStringSet on the whole-genome protein sequences to pre-process the primary transcript only input data. Without this step, the number of duplicate genes will be overrepresented. In the Figure 6 of Wang et al., three closely related Arabidopsis taxa were compared in terms of different gene duplication modes. In our investigation of this figure, we found that the data for *Arabidopsis arenosa* (23,097 protein genes) and *Arabidopsis suecica* (63,469 protein genes) were based on proteins from primary transcripts, but this was not the case for *Arabidopsis thaliana* (48,265 protein genes).

Here, we show how using a primary protein filtering step can dramatically impact the results of this previous analysis by Wang et al. (Figure 1). Indeed, by following the same MCScanX steps in the protocol, we found as many as 25,776 protein genes in singleton duplication after employing primary protein filtering, compared to only 3,086 protein genes when not filtering. Similarly, there are 11,948 protein genes in dispersed duplication when filtering vs. 7,297 protein when not. Finally, there are only 2,216 protein genes in tandem duplication after the primary protein filtering compared to 30,101 protein genes without filtering. Strikingly, it appears that tandem duplications are less prevalent in *A. thaliana* than *A. arenosa*, and singleton duplication composes an overwhelming proportion of gene duplications in *A. thaliana* compared to other two species.

**Figure 1.**
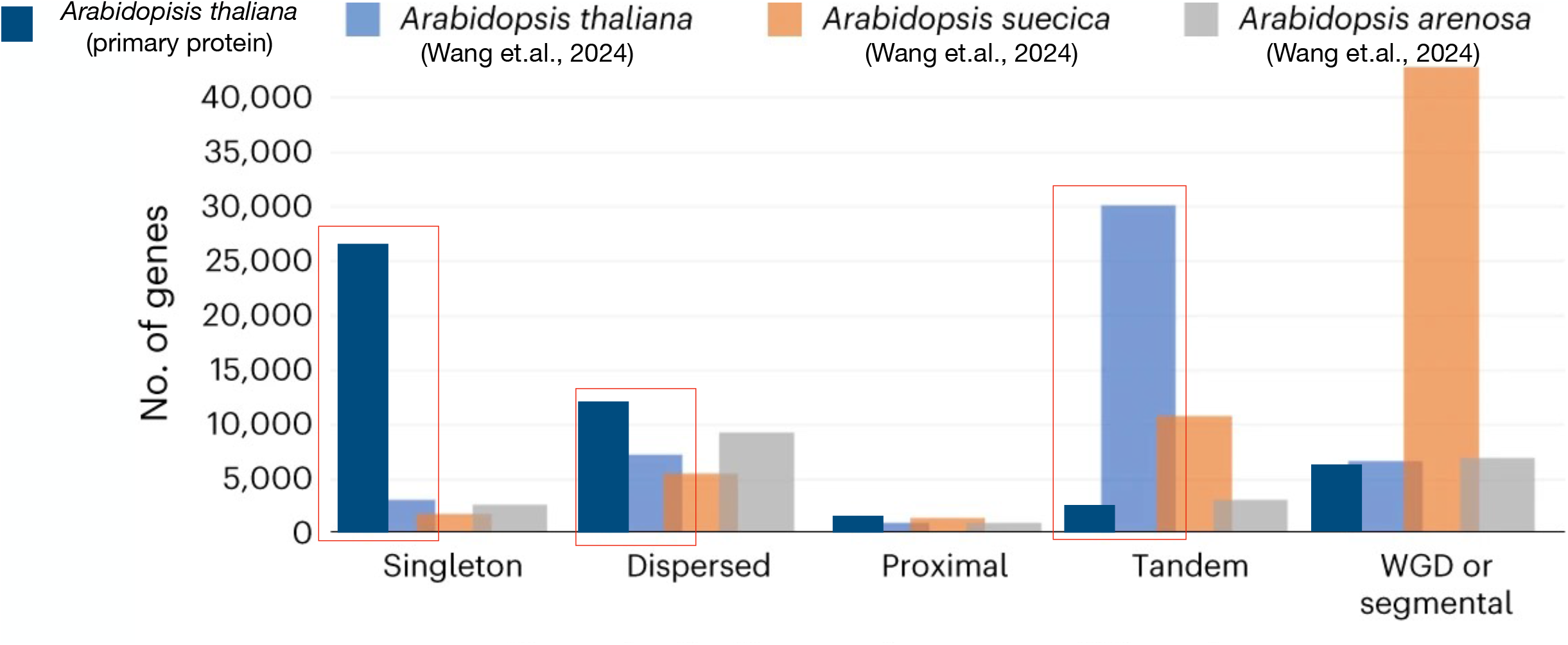
Comparison of gene duplication modes among closely related Arabidopsis taxa. The figure was adapted from Figure 6 of the protocol by Wang et al. (2024) (Wang et al., 2024). *A. thaliana* after primary filtering was in dark blue and the differences between the rest of species (*A. arenosa* in grey and *A. suecica* in brown, and *A*.*thaliana* without filtering in light blue) in the bar graph have been highlight in red.

We found the labelling of the X-axis in Figure 5 of Wang et al. is inaccurate; the authors used a different version of the *A. thaliana* genome (Araport11) as compared to the one demonstrated in the protocol (TAIR 10). Considering that the Araport11 version only has a few more protein-coding genes than the TAIR 10 version, we used the TAIR10 of *A. thaliana* to ensure the data consistency throughout the protocol. As shown in Figure 2, we chose *A. thaliana* and *Medicago truncatula* for the visualization of the synonymous (Ks) substitution rate distributions of colinear genes before and after primary protein filtering.

**Figure 2.**
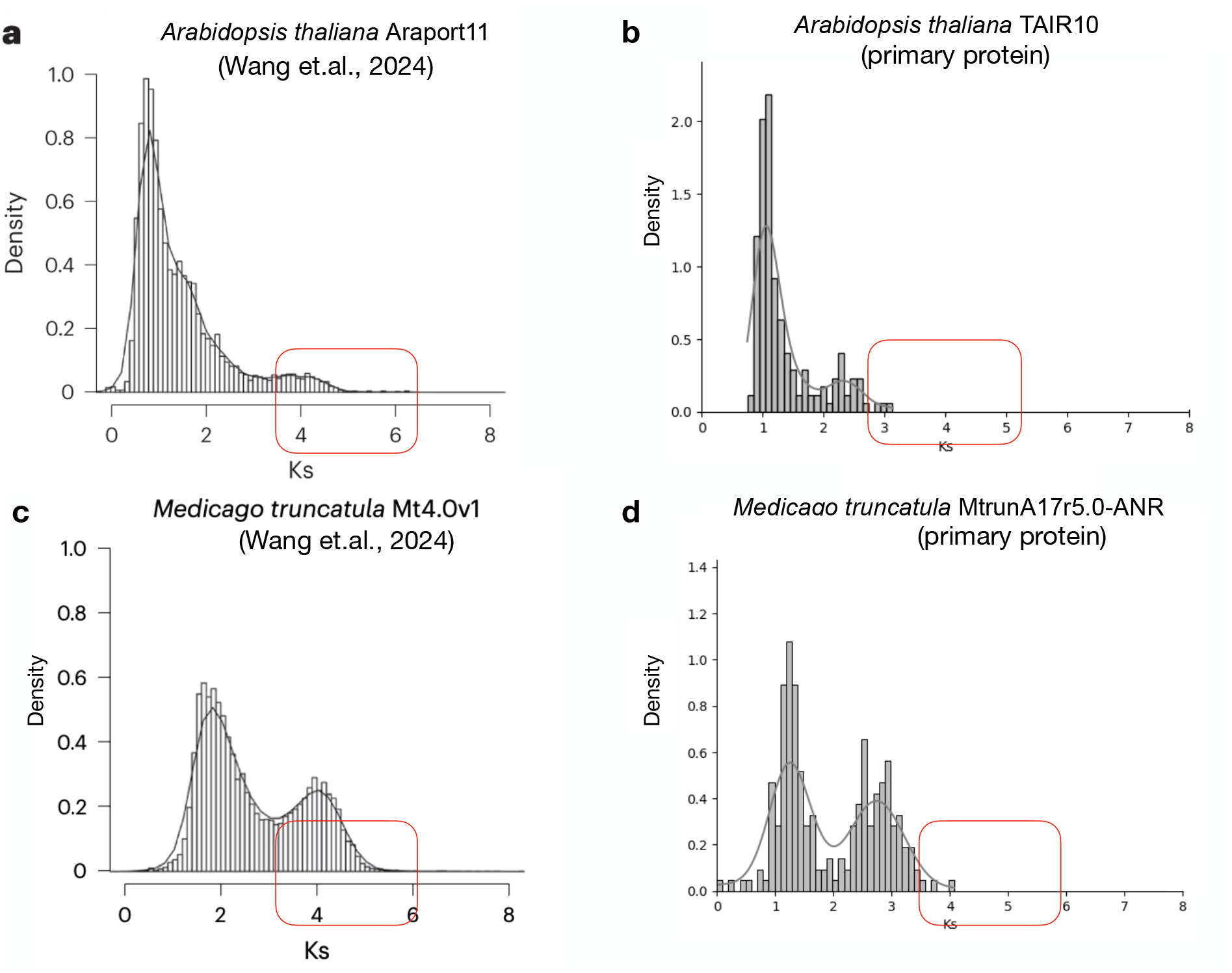
Visualization of the synonymous (Ks) substitution rate distributions of colinear genes in different genomes (*A*.*thaliana* and *M. truncatula*) Note: The figure a and c were adapted from Figure 5 of the protocol (Wang et al., 2024). The difference of the X-axis is highlighted in red before and after the primary protein filtering. The *add_Ka_and_Ks_to_collinearity_yn00. pl* program from the protocol was applied to add evolutionary rates to each ‘. collinearity’ file, and its Ks distribution was visualized by custom python script from the GitHub: https://github.com/zx0223winner/MCScanX_Assistant.

The density curve is similar in *M. truncatula* (Figure 2c and 2d) with the exception that most of the Ks distribution values are smaller than four in our analysis compared to five in the protocol by Wang et al. Similarly, the Ks distribution values are mostly smaller than three in our analysis of *A. thaliana* (Figure 1b), compared to four in the protocol (Figure 1a). This is consistent with the *A. thaliana* result without any primary protein filtering (Supplementary Figure 1). Here, the density curve is sharper (Figure 1a) for the *A. thaliana* data without primary protein filtering compared to the one that was filtered (Figure 1b). This is because we could not find the R script - ‘visualize_Ks_distributions.R’ demonstrated at Step 26-A-(iv); instead, we built the code from scratch to generate the similar density histogram plots to reflect the patterns. Density usually refers to the relative frequency of data points within a specific bin, scaled by the bin’s width. In terms of the density difference in Y-axis, we do not know if Wang et al. applied any normalization strategy, but the density at Y-axis can be a high value when the histogram plot was drawn on narrow bins with high concentration of data points. This is consistent with the supplementary plot (Supplementary Figure 2) when we add a rug plot which shows every single data on the X-axis turning out those pillars with high values are in this case. Given all these, we observed the pattern of two peaks in the Ks distributions still stands out as indicators (before and after primary protein filtering) for the potential large-scale gene duplication events.

We are not questioning the reliability of running the MCScanX algorithms but do want to raise awareness of the potential challenges when preparing the input files in Part 2 (Steps 7–20) and also want to highlight potential issues when using the protocol. Here are some other useful tips, as the authors detailed at step 11-13 and the troubleshooting section on the generation of correct ‘.gff’ file, we found it is better to offer more specific backup scripts options similar to the ‘mkGFF3.pl’ program in MCScanX_protocol package. This is because the downloaded ‘.gff’ can have different formats and it is important to convert it to the one MCScanX can read. We have found that the ‘gff2bed’ script from BEDOPS v2.4.41 (Neph et al., 2012) and the custom processing script on the ‘XX_feature_table.txt’ (GitHub: https://github.com/zx0223winner/MCScanX_Assistant) can help yield the ‘.gff’ file for MCScanX. In terms of generating the ‘.blast’ file at Step 14-20, we found it is not efficient to prepare the ‘runBLASTP.sh’, especially when all-against-all BLASTP are needed for each reciprocal genome pair. We have prepared a custom script from the same GitHub link above to iterate the genome all-against-all BLASTP processing which greatly increases the preparation step. Overall, MCScanX is a very useful tool for efficiently identifying colinear blocks and downstream evolutionary analysis, but additional work is needed for preparing the input data and running the tool.

## Supporting information

Supplementary data for methodological details

## Acknowledgments

XZ conceptualized the project and analyzed associated data. The manuscript was written by XZ. All authors read, revised, and approved the final manuscript for peer review.

## Availability and implementation

The codes in protocol use SnakeMake pipeline and Conda to run and install dependencies. The distribution version is available online at GitHub: https://github.com/zx0223winner/MCScanX_Assistant and the archived version at Zenodo is https://doi.org/10.5281/zenodo.16423847. Methodological details can be found via the link: https://github.com/zx0223winner/MCScanX_Assistant/blob/main/docs/Usage.md.

## Supplementary materials

**Supplementary Figure S1.**
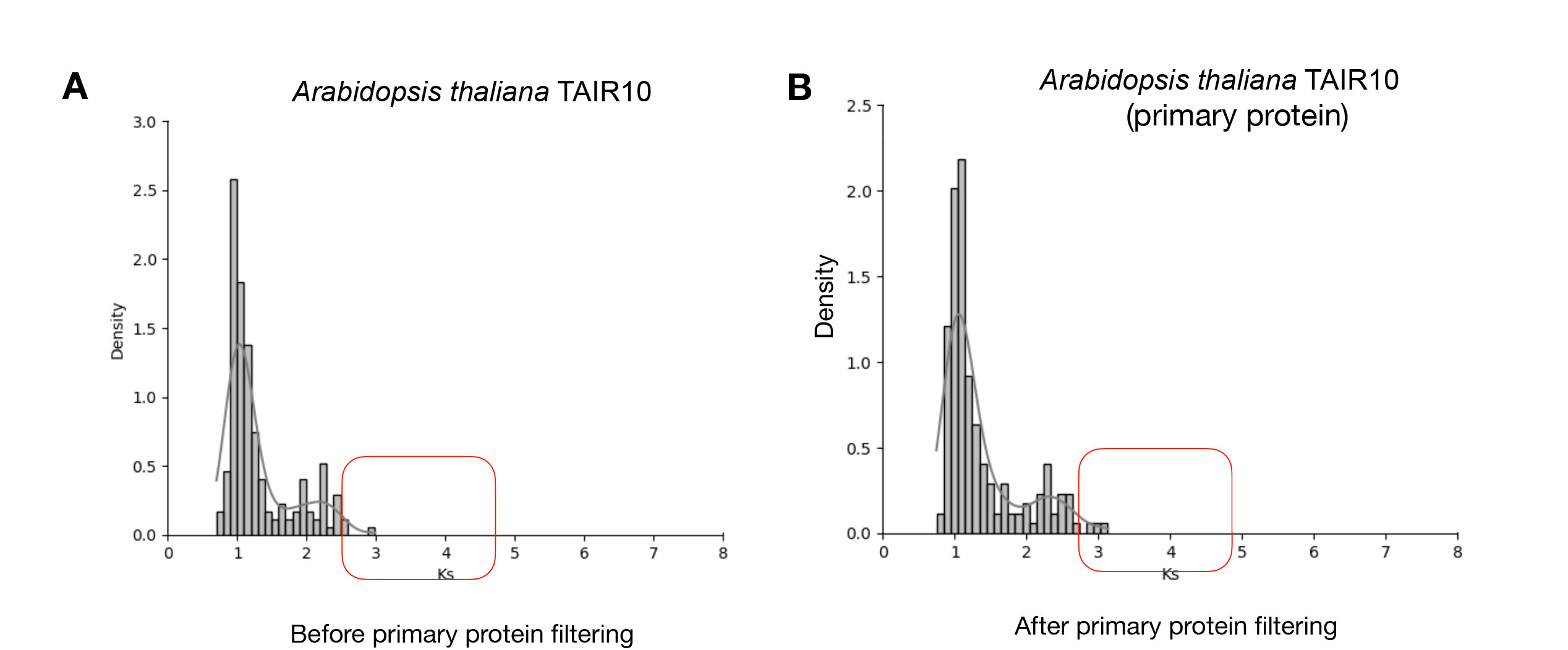
The synonymous (Ks) substitution rate distributions of colinear genes in *A. thaliana* without any primary protein filtering.

**Supplementary Figure S2.**
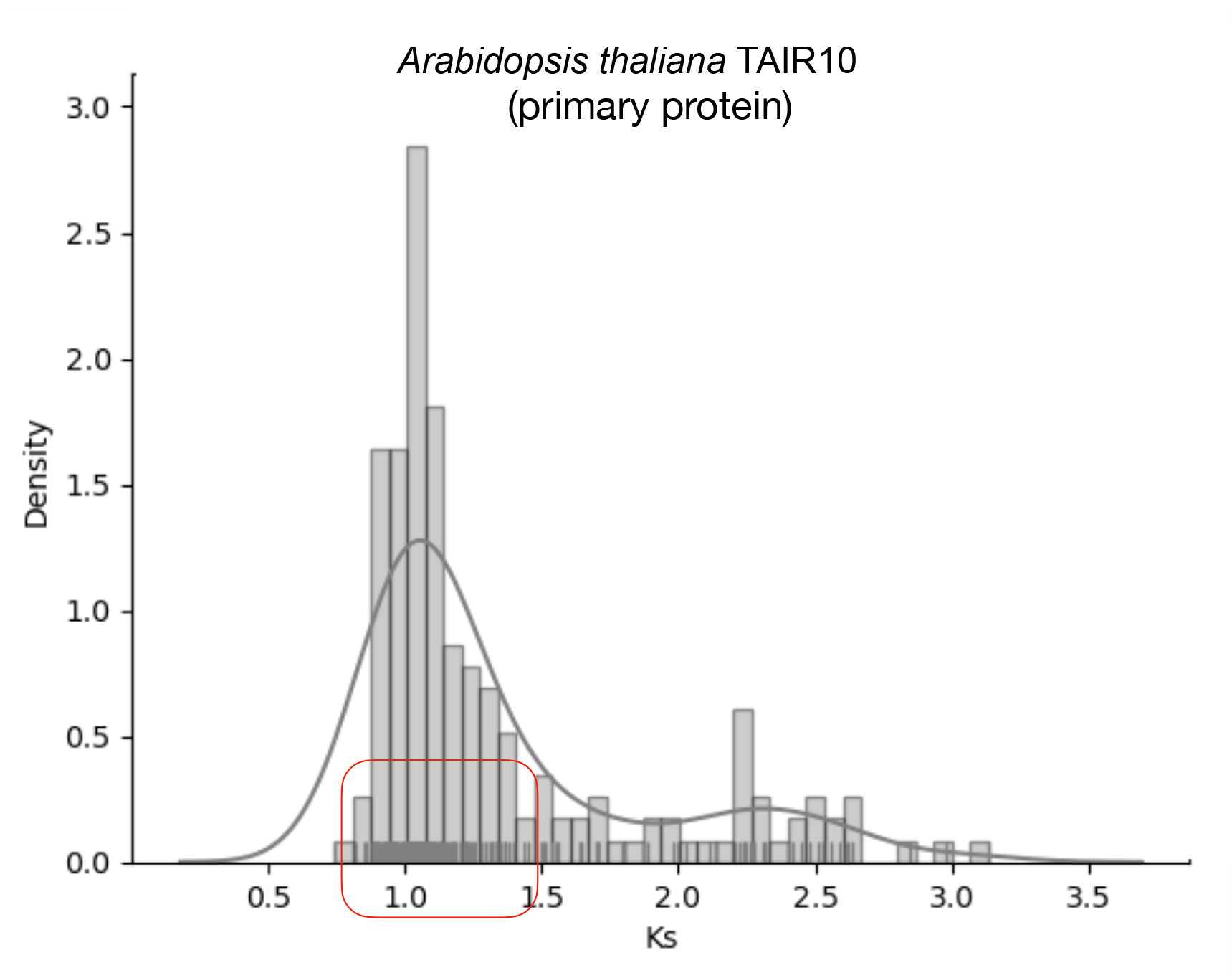
The synonymous (Ks) substitution rate distributions of colinear genes in *A. thaliana* after primary protein filtering together with a rug plot indicating the density intensity.

