## Supplementary data for methodological details for "How to prepare the input data and run MCScanX efficiently?"

### MCScanX_Assistant: Supplementary materials

#### 📈 MCScanX_Assistant workflow.

- Part 1: Snakefile_input_preparing
- (1)Prepare the SnakeMake config file which contains the species name, NCBI genomic assembly ID and other input file directories.
- (2)Prepare the '.gff' and diamond BLASTP '.blast' files for MCScanX protocol.
- (3)Provide alternative options for preparing the '.gff' files and iterate the all-vs-all blastp on all genome pairs

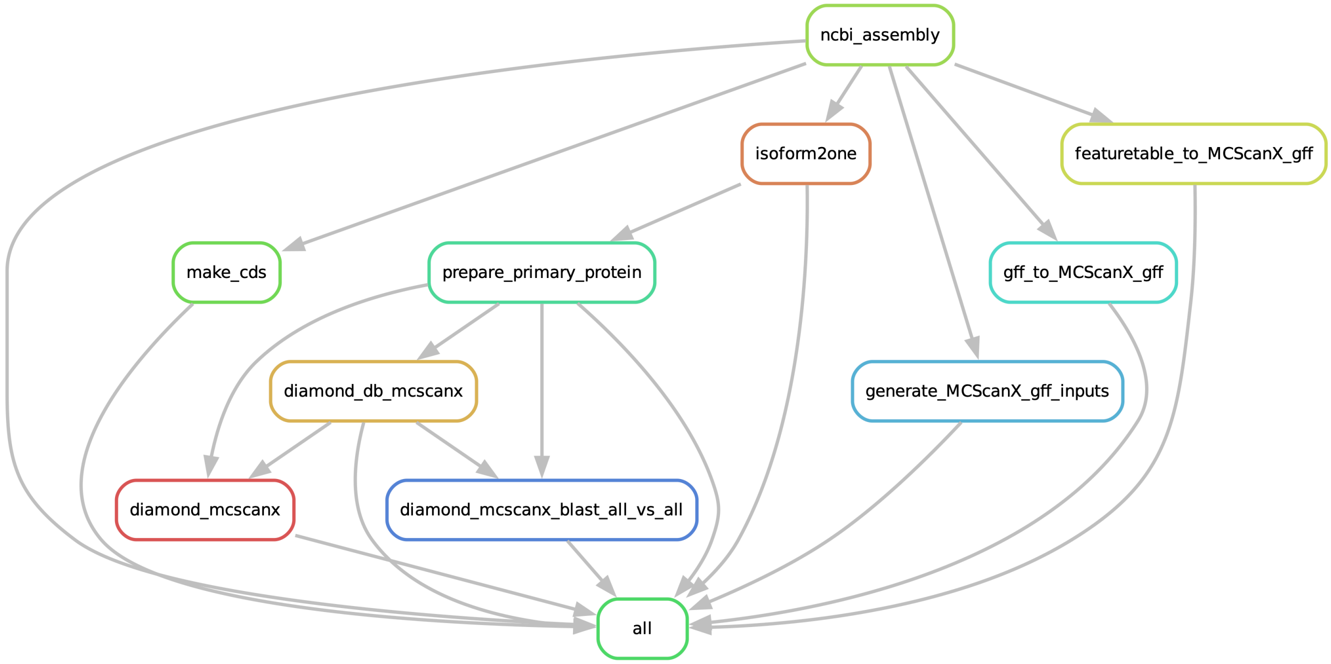


- Part 2: Snakefile_Ks_distribution_plot
- (1)Calculating the kaks via the perl script
- (2)Display the distribution of Ks peaks
- (3)create the density plot for the Ks values

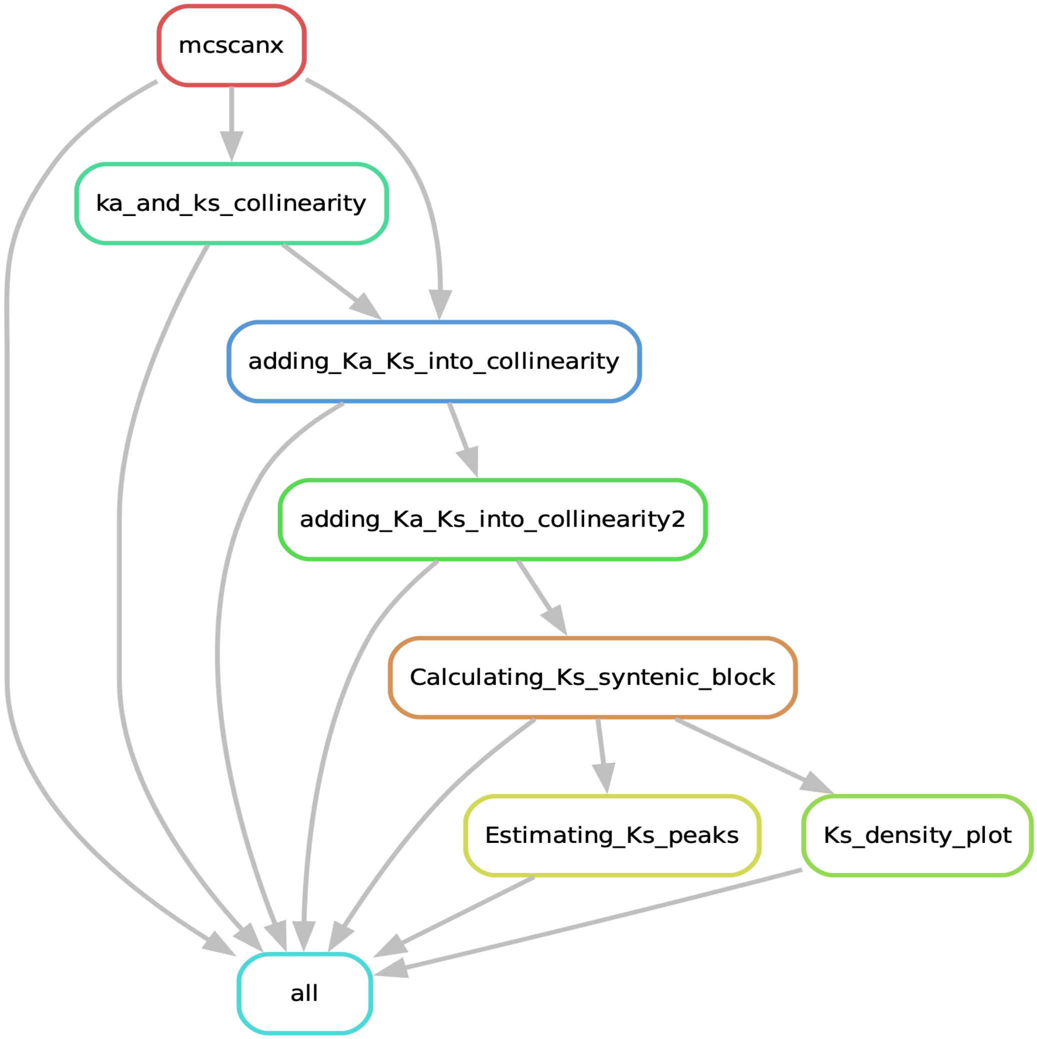


#### 📈 MCScanX_Assistant workflow.

- Part 3: Snakefile_MCScanX_6species
- Reproduce what the protocol did for the 6 species all together
- This is useful for users who want to apply multiple different species in MCScanX running.

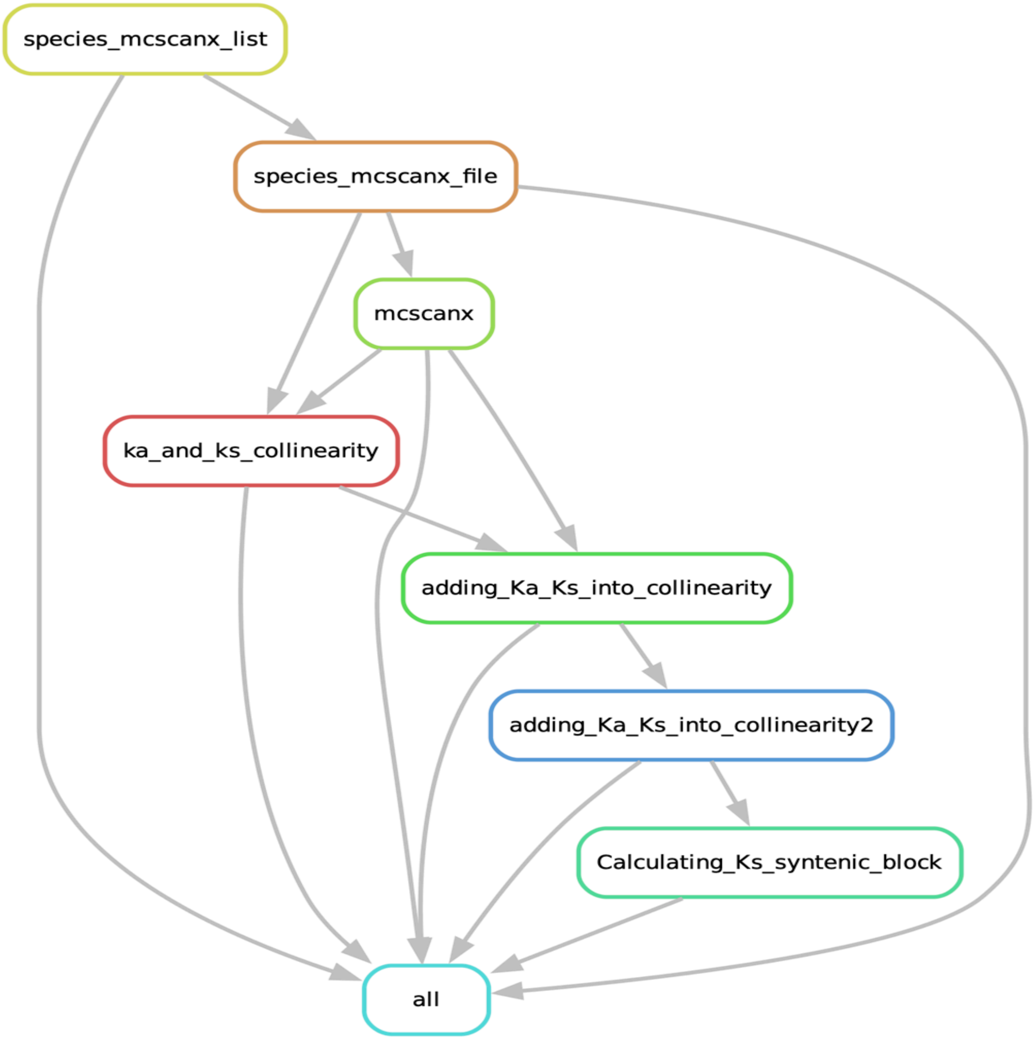


#### 📋Supplementary Text : Usage of SnakeMake pipeline.

Contents:


- Text S1. Introduction for the config.yaml file;
- Text S2. Download and preprocess the NCBI assemblies (Snakefile_Input_preparing);
- Text S3. Visualization of the Ks distribution plots (Snakefile_Ks_distribution_plot);
- Text S4. Reproduce the pre-processing of 6 species in the protocol (Snakefile_MCScanX_6species).


#### Text S1. Config.yaml file

You will need to edit the config.yaml file for your own usage. An example config.yaml has been provided to test the pipeline.

> ⚠ **Warning**
>
> feel free to modify the species name with yours, keep the similiar input file format.

```yaml
ncbi_assemblies:
  - GCA_016771965.1
  - GCA_019202805.1
  - GCA_905216605.1
  - GCA_911865555.2
  - GCF_000001735.4
  - GCF_000695525.1

names:
  - "Bcarinata"
  - "Asuecica"
  - "Aarenosa"
  - "Tarvense"
  - "Athaliana"
  - "Boleracea"


ks_test_name:
  - "6species"

species_name:
  - Athaliana
 # - Mtruncatula

ncbi_genomes:
    Bcarinata:
        ncbi_assembly: "data/ncbi_download/GCA_016771965.1.zip"
        assembly_id: "GCA_016771965.1"
        feature_table: "data/ncbi_download/GCA_016771965.1_ASM1677196
        species: Bcarinata

    Asuecica:
        ncbi_assembly: "data/ncbi_download/GCA_019202805.1.zip"
        assembly_id: "GCA_019202805.1"
        feature_table: "data/ncbi_download/GCA_019202805.1_ASM1920280
        species: Asuecica


    Aarenosa:
        ncbi_assembly: "data/ncbi_download/GCA_905216605.1.zip"
        assembly_id: "GCA_905216605.1"
        feature_table: "data/ncbi_download/GCA_905216605.1_AARE701a_f
        species: Aarenosa
```

```
    Tarvense:
        ncbi_assembly: "data/ncbi_download/GCA_911865555.2.zip"
        assembly_id: "GCA_911865555.2"
        feature_table: "data/ncbi_download/GCA_911865555.2_T_arvense_
        species: Tarvense


    Athaliana:
        ncbi_assembly: "data/ncbi_download/GCF_000001735.4.zip"
        assembly_id: "GCF_000001735.4"
        feature_table: "data/ncbi_download/GCF_000001735.4_TAIR10.1_f
        species: Athaliana


    Boleracea:
        ncbi_assembly: "data/ncbi_download/GCF_000695525.1.zip"
        assembly_id: "GCF_000695525.1"
        feature_table: "data/ncbi_download/GCF_000695525.1_BOL_featur
        species: Boleracea


    Rsativus:
        ncbi_assembly: "data/ncbi_download/GCF_000801105.2.zip"
        assembly_id: "GCF_000801105.2"
        feature_table: "data/ncbi_download/GCF_000801105.2_ASM80110v3
        species: Rsativus


    Bnapus:
        ncbi_assembly: "data/ncbi_download/GCF_020379485.1.zip"
        assembly_id: "GCF_020379485.1"
        feature_table: "data/ncbi_download/GCF_020379485.1_Da-Ae_feat
        species: Bnapus

    # Mtruncatula:
    #     ncbi_assembly: "data/ncbi_download/GCF_003473485.1.zip"
    #     assembly_id: "GCF_003473485.1"
    #     feature_table: "data/ncbi_download/GCF_003473485.1_MtrunA17
    #     outgroups: "Arabidopsis_thaliana"
    #     species: Mtruncatula

MCScanX_protocol:
 - "/scripts/MCScanX_protocol"
```

#### Text S2. Snakefile_Input_preparing

#### Download NCBI assemblies

`Purpose`: This rule provides a convenient way to download the standard input files from NCBI.

> ⓘ Note
>
> To avoid repeatly download the ".zip" files with the example file we provided ('MCScanX_Assistant_data.tar.gz'), we commented the rule in the snakefile.

`scripts`:

```
mkdir -p {params.dir};\
curl -OJX \
GET "{params.link}"; \
mv {params.file} {params.dir} \
```

`Output`: data/ncbi_download/GCF_000001735.4.zip

```
# standard input files from NCBI
XX.genomic.gff
XX.protein.faa
XX.cds_from_genomic.fna
```

> ⓘ Note
>
> Optional: To download extra ncbi assembly 'XX.zip' from NCBI, users can substitue the ncbi_assembly id (e.g., GCF_000001735.4) with yours in the command below:

```
curl -OJX GET
"https://api.ncbi.nlm.nih.gov/datasets/v2alpha/genome/accession/GCF_0
include_annotation_type=GENOME_FASTA,GENOME_GFF,RNA_FASTA,CDS_FASTA,P
```

#### Preprocessing the naming of the NCBI assemblies

`Purpose`: Rename the NCBI genomic assembly to the format which mcscanx and Dupgen-finder can take.

`scripts`:

```
mkdir -p {params.dir2}{params.species_name}; \
unzip {params.dir1}{params.assembly_id}.zip -d {params.dir1}
{params.species_name}; \
sleep 5s; \
cp {params.dir1}
{params.species_name}/ncbi_dataset/data/{params.assembly_id}/cds_from
{params.dir2}
{params.species_name}/{params.species_name}_cds_from_genomic.fna;
\
cp {params.dir1}
{params.species_name}/ncbi_dataset/data/{params.assembly_id}/genomic.
{params.dir2}
{params.species_name}/{params.species_name}_genomic.gff; \
cp {params.dir1}
{params.species_name}/ncbi_dataset/data/{params.assembly_id}/protein.
{params.dir2}
{params.species_name}/{params.species_name}_protein.faa; \
rm -r {params.dir1}{params.species_name} \
```

`Output` : "data/ncbi/Athaliana/Athaliana_genomic.gff" ; "data/ncbi/Athaliana/Athaliana_protein.faa"; "data/ncbi/Athaliana/Athaliana_cds_from_genomic.fna".

#### Preprocessing the gff file (default)

`Purpose` : Create a MCScanX_gff from the gff3 which can be recognized by McscanX

> ⓘ Note
>
> Since the required input .gff file for mcscanx is nether gff3 nor bed file format, for simplity, call it MCScanX_gff file

> Tip
>
> If the mkGFF3.pl does not work on your gff3 file due to the format of naming, there are other ways/options to generate the MCScanX_gff, check the next rules and substitue the gff with the one works.

> ⚠ Warning
>
> The mkGFF3.pl was adopted from MCScanX_protocol which is not exactly same (Wang, Yupeng, et al. Nature Protocols 19.7 (2024): 2206-2229.)

`scripts`:

```
mkdir -p {params.dir};\
curl -OJX \
GET "{params.link}"; \
mv {params.file} {params.dir} \
```

`Input`: data/ncbi/Athaliana/Athaliana_genomic.gff

```
##gff-version 3
#!gff-spec-version 1.21
#!processor NCBI annotwriter
#!genome-build TAIR10.1
#!genome-build-accession NCBI_Assembly:GCF_000001735.4
#!annotation-source TAIR and Araport
##sequence-region NC_003070.9 1 30427671
##species
https://www.ncbi.nlm.nih.gov/Taxonomy/Browser/wwwtax.cgi?id=3702
NC_003070.9     RefSeq  region  1       30427671        .       +
.
ID=NC_003070.9:1..30427671;Dbxref=taxon:3702;Name=1;chromosome=1;ecot
DNA
```

`Output`: data/intermediateData/Athaliana/Athaliana.gff

```
#gff
Athaliana1      NP_171609.1     3760    5630
Athaliana1      NP_001318899.1  6915    8666
Athaliana1      NP_001321777.1  6915    8442
Athaliana1      NP_001321775.1  6915    8442
Athaliana1      NP_001321776.1  6915    8419
```

#### Preprocessing the gff (option one)

`Purpose`: This rule uses the gff2bed tool to convert gff to bed for easier parsing (the MCScanX_gff file)

> ⚠ Warning

> To run this rule, users will need uncomment the lines in the workflow/snakefile_part1 file

```
        #       expand("data/intermediateData/{name}/{name}.gff-option_one",
        #               name = config['names']),
```

`scripts`:

```
cat {input.gff} \
| grep -v '^#' \
| awk '$3 == "gene"' \
| gff2bed \
| awk 'BEGIN {{OFS="\t"}} {{print $1,$4,$2,$3}}' \
> {output.MCScanXgff} \
```

`Output`: data/intermediateData/Athaliana/Athaliana.gff-option_one

#### Preprocessing the gff (option Two)

`Purpose`: This rule can make use of the XX.feature_table.txt from NCBI to generate the MCScanX_gff for McScanX as input file.

```
# Example of the XX.feature_table.txt:
link:
https://ftp.ncbi.nlm.nih.gov/genomes/all/GCF/000/001/735/GCF_00000173
```

`scripts`:

```
sed 1d {input.feature_table} \
|grep 'mRNA' \
|awk -F'\t' '$13!=""{{print $7"\t"$13"\t"$8"\t"$9}}' \
> {output.MCScanXgff} \
```

`Output`: data/intermediateData/Athaliana/Athaliana.gff-option_two

> ⚠ Warning
>
> To run this rule, user will need uncommenting the lines in the snakefile_part1

```
        # Other ways to yield the MCScanXgff: featuretable_to_MCScanXgff
        #       expand("data/intermediateData/{name}/{name}.gff-option_two",
        #               name = config['names']),
```

#### Prepare the cds file for calculating the Ka/ks ratio

`Purpose`: This rule is preprocessing step for running the McScanX with input data from genomic cds

> ⚠ **Warning**
>
> The mkCD.pl was adopted from MCScanX_protocol which is not exactly the same (Wang, Yupeng, et al. Nature Protocols 19.7 (2024): 2206-2229.)

`scripts`:

```
perl {params.dir2}/mkCD.pl {params.dir3} {params.species_name} \
```

`Output`: data/intermediateData/Athaliana/Athaliana.cds

```
>NP_171609.1
ATGGAGGATCAAGTTGGGTTTGGGTTCCGTCCGAACGACGAGGAGCTCGTTGGTCACTATCTCCGTAAC
AAACACTAGCCGCGACGTTGAAGTAGCCATCAGCGAGGTCAACATCTGTAGCTACGATCCTTGGAACTT
```

#### Prepare the primary protein for the input file

`Purpose`: This rule is the preprocessing step for extracting the longest transcript encoding for each gene, and use the primary protein for the rest of analysis.

> ⓘ **Note**
>
> Due to alternative splicing, the mRNA isoform/transcript can have different lengths, which encoding the protein product with different ID but from same gene. This step is to minimize the misprediction of gene duplicates for those proteins encoded by alternative splicing transcitps having similiar fucntional domains.

`scripts`:

```
        python3 {params.dir1}/isoform2one.py {input.feature_table}
{output}; \
        awk '{{print $1}}' {input.protein} \
```

`Output` : "data/ncbi/{name}_primary/{name}_protein.list", "data/ncbi/{name}_primary/{name}_protein.faa",

```
# Athaliana_protein.list
NP_171609.1
NP_001321775.1
NP_171611.1
NP_171612.1
NP_171613.1
NP_001320628.
```

> **Note**
>
> There are rare cases for NCBI without feature table to download (e.g., https://ftp.ncbi.nlm.nih.gov/genomes/all/GCF/000/001/735/GCF_000001735.4_TAIR10.1/GCF_000001735.4_TAIR10.1_feature_table.txt.gz). Users can prepare primary protein gene list - "XX_protein.list" from NCBI website manually, For example, the proteins column for Athaliana: https://www.ncbi.nlm.nih.gov/datasets/gene/GCF_000001735.4/?gene_type=protein-coding

#### Diamond_db_mcscanx

`Purpose` : This rule builds diamond database for blasting the protein sequence

`scripts` :

```
        mkdir -p {params.dir1}; \
        diamond makedb \
                --in {params.protein} \
                -d {params.db_name_dir} \
```

`Output` : data/ncbiDB/Athaliana.dmnd

#### Diamond_blast_mcscanx

`Purpose`: This rule runs diamond blastp for the protein sequences against themselves (blastp all vs all)

> ⓘ Note
>
> --max-target-seqs parameter will impact how many candidate duplicates will be detected `scripts`:

```
                 diamond blastp \
                 -d {params.db_name_dir} \
                 -q {params.protein} \
                 -o {output} \
                 -e 1e-10 \
                 -f 6 \
                 -p {threads} \
                 --sensitive \
                 --max-target-seqs 5 \
```

`Output`: data/intermediateData/Athaliana/Athaliana.blast

```
NP_001030613.1  NP_001030613.1  100     596     0       0       1
596     1       596     0.0     1155
NP_001030613.1  NP_001327195.1  100     583     0       0       1
583     47      629     0.0     1132
NP_001030613.1  NP_186759.2     100     583     0       0       1
583     1       583     0.0     1132
```

#### diamond_mcscanx_blast_all_vs_all

`Purpose`: This rule can iterate the all-vs-all blastp on all genome pairs. For six species (6^2=36 times)

> ⓘ Note
>
> --max-target-seqs parameter will impact how many candidate duplicates will be detected `scripts`:

```
diamond blastp \
-d {params.db_name_dir} \
-q {params.protein} \
-o {output} \
-e 1e-10 \
-f 6 \
-p {threads} \
--sensitive \
--max-target-seqs 5 \
```

`Output`: "data/intermediateData/{name_a}-{name_b}_mcscanx/{name_a}-{name_b}.blast"

### Text S3. Snakefile_Ks_distribution_plot

#### Run the MCScanX

`Purpose`: This is the major script to run the MCScanX scripts with the previous prapered '.gff' and '.blast' files.

`scripts`:

```
export PATH=$PATH:{params.dir2}; \
chmod +x {params.dir2}/MCScanX; \
cp -r {params.dir} {params.dir2}; \
./{params.dir2}/MCScanX {params.dir1}/{params.dir_name} \
```

`Output`: "scripts/MCScanX/{species_name}/{species_name}.collinearity"

> ⓘ Note
>
> If mcscanx has no results, solution is to put .blast and .gff file inside MCScanX folder without creating a separate folder, the above rule can do this.

#### Calculating ka_and_ks values from the duplicates pairs

`Purpose`: This rule can run the PAML package (Yang, Ziheng.Molecular biology and evolution 24.8 (2007): 1586-1591.) to calculate the kaks for the gene pairs.

> ⚠ Warning

> The add_ka_and_ks_to_collinearity_Yn00.pl was adopted from DupGen_finder which is not exactly same (Qiao, Xin, et al. Genome biology 20 (2019): 1-23; Wang, Yupeng, et al. Nucleic acids research 40.7 (2012): e49-e49).
> https://github.com/qiao-xin/Scripts_for_GB/tree/master/identify_Ks_peaks_by_fitting_GMM

`scripts`:

```
        cp {params.dir1}/{params.species_name}.cds
{params.dir5}/{params.species_name}.cds; \
        sleep 30s; \
        perl {params.dir4}/add_ka_and_ks_to_collinearity_Yn00.pl \
        -i {input} \
        -d {params.dir5}/{params.species_name}.cds \
        -o {output} \
```

`Output`: data/DupGen_finder/Athaliana_result/Athaliana.kaks

```
NP_001321164.1  NP_001185394.1  0.2543  0.7844  0.3242    2e-73
NP_001322884.1  NP_001320573.1  0.2128  0.9219  0.2308        0
NP_564051.1     NP_001323057.1  0.1533  0.8397  0.1826    4e-62
NP_173281.1     NP_565075.1     0.1210  0.8860  0.1365        0
NP_564052.1     NP_001322804.1  0.1003  0.5209  0.1926   2e-270
NP_173285.2     NP_001322762.1  0.0908  0.9425  0.0963   1e-281
NP_173286.2     NP_683494.2     0.2178  1.3273  0.1641   4e-131
NP_173289.1     NP_177545.1     0.0523  0.9223  0.0567   3e-147
```

> **Note**
>
> Due to the perl script: add_ka_and_ks_to_collinearity_Yn00.pl, which may have some temporary files left in main dir, which can be safefly removed.

```
#       rm -p *.aln; \
#       rm -p *.cds; \
#       rm -p *.dnd; \
#       rm -p *.pro; \
```

#### Adding_Ka_Ks_into_collinearity

`Purpose`: This rule is to preprocess the XX.collinearity and XX.kaks file for the next step

`scripts`:

```
        mkdir -p {params.dir}; \
        cp {input.col}
{params.dir}/{params.species_name}.collinearity; \
        awk -F'\t' '{{print $2"\t"$3"\t"$5"\t"$6"\t"$7"\t"$4}}'
{input.kaks} \
|grep -v -e '^[[:space:]]*$' \
> {output[0]}
```

`Output`: data/DupGen_finder/Athaliana_result_kaks/Athaliana.collinearity

```
############### Parameters ###############
# MATCH_SCORE: 50
# MATCH_SIZE: 5
# GAP_PENALTY: -1
# OVERLAP_WINDOW: 5
# E_VALUE: 1e-05
# MAX GAPS: 25
############### Statistics ###############
# Number of collinear genes: 6451, Percentage: 13.40
# Number of all genes: 48147
##########################################
## Alignment 0: score=4086.0 e_value=0 N=91 Athaliana1&Athaliana1
plus
  0-  0:        NP_001321164.1  NP_001185394.1    2e-73
  0-  1:        NP_001322884.1  NP_001320573.1        0
  0-  2:        NP_564051.1     NP_001323057.1    4e-62
```

#### Adding_Ka_Ks_into_collinearity2

`Purpose`: This rule can add Ka, Ks, Ka/Ks values into Athaliana.collinearity by using Athaliana.kaks as input, and produce one output file: Athaliana.collinearity.kaks

> ⚠ Warning

> The add_ka_ks_to_collinearity_file.pl was adopted from DupGen_finder which is not exactly the same (Qiao, Xin, et al. Genome biology 20 (2019): 1-23; Wang, Yupeng, et al. Nucleic acids research 40.7 (2012): e49-e49). [https://github.com/qiao-xin/Scripts_for_GB/tree/master/identify_Ks_peaks_by_fitting_GMM](https://github.com/qiao-xin/Scripts_for_GB/tree/master/identify_Ks_peaks_by_fitting_GMM)

`scripts`:

```
perl {params.dir1}/add_ka_ks_to_collinearity_file.pl
{params.dir2}/{params.species_name} \
```

`Output`: data/DupGen_finder/Athaliana_result_kaks/Athaliana.collinearity.kaks

```
############### Parameters ###############
# MATCH_SCORE: 50
# MATCH_SIZE: 5
# GAP_PENALTY: -1
# OVERLAP_WINDOW: 5
# E_VALUE: 1e-05
# MAX GAPS: 25
############### Statistics ###############
# Number of collinear genes: 6451, Percentage: 13.40
# Number of all genes: 48147
##########################################
## Alignment 0: score=4086.0 e_value=0 N=91 Athaliana1&Athaliana1
plus
  0-  0:        NP_001321164.1  NP_001185394.1    2e-73 0.2543
0.7844  0.3242
  0-  1:        NP_001322884.1  NP_001320573.1        0 0.2128
0.9219  0.2308
  0-  2:        NP_564051.1     NP_001323057.1    4e-62 0.1533
0.8397  0.1826
  0-  3:        NP_173281.1     NP_565075.1           0 0.1210
0.8860  0.1365
```

#### Calculating_Ks_syntenic_block

`Purpose`: This rule produces one output file: Athaliana.synteny.blocks.ks.info, which contains average Ks values for gene pairs contained in each syntenic block.

> ⚠ Warning

> The compute_ks_for_synteny_blocks.pl was adopted from DupGen_finder which is not exactly same (Qiao, Xin, et al. Genome biology 20 (2019): 1-23; Wang, Yupeng, et al. Nucleic acids research 40.7 (2012): e49-e49). https://github.com/qiao-xin/Scripts_for_GB/tree/master/identify_Ks_peaks_by_fitting_GMM

`scripts`:

```
        perl {params.dir1}/compute_ks_for_synteny_blocks.pl
{input}; \
        cp {params.species_name}.synteny.blocks.ks.info {output};
\
        rm {params.species_name}.synteny.blocks.ks.info \
```

`Output`: data/DupGen_finder/Athaliana_result_kaks/Athaliana.synteny.blocks.ks.info

```
Blocks ID       Location        Block Size      Average Ks      e-
value   Score   Orientation
Alignment169    Athaliana5&Athaliana5   16      0.94783125
2.9e-41 719.0   plus
Alignment105    Athaliana2&Athaliana5   10      2.41932 8.7e-21
430.0   minus
Alignment159    Athaliana4&Athaliana5   6       1.43243333333333
8.6e-09 262.0   plus
Alignment9      Athaliana1&Athaliana1   14      1.66228571428571
8.2e-34 640.0   plus
Alignment148    Athaliana3&Athaliana5   6       0.922966666666667
0       264.0   minus
Alignment119    Athaliana3&Athaliana5   95      0.920378494623656
0       4241.0  plus
```

#### Estimating Ks peaks from Ks distribution

`Purpose`: This rule can create the Ks distribution of Ks values of syntenic blocks within the genome

> ⚠ Warning
>
> The plot_syntenic_blocks_ks_distri.py was adopted from DupGen_finder which is not exactly same (Qiao, Xin, et al. Genome biology 20 (2019): 1-23; Wang, Yupeng, et al. Nucleic acids research 40.7 (2012): e49-e49). https://github.com/qiao-xin/Scripts_for_GB/tree/master/identify_Ks_peaks_by_fitting_GMM

> ℹ️ Note
>
> The parameter 'Components' can indicate the number of the mixture components, which represents the number of Ks peak.

`scripts`:

```
perl {params.dir1}/plot_syntenic_blocks_ks_distri.py {input}
{params.components} {params.dir2}/{params.species_name} \
```

`Output`:
data/DupGen_finder/Athaliana_result_kaks/Athaliana.synteny.blocks.ks.distri.pdf

#### Ks_density_plot

`Purpose`: This rule can create the density plot of Ks distribution

> ℹ️ Note
>
> The parameter 'Components' can indicate the number of the mixture components, which represents the number of Ks peak.

`scripts`:

```
python {params.dir1}/plot_density_ks_distri.py {input}
{params.dir2}/{params.species_name}.synteny.density.ks.distri.pdf
{params.species_name}\
```

`Output`:
data/DupGen_finder/Athaliana_result_kaks/Athaliana.synteny.blocks.ks.distri.pdf

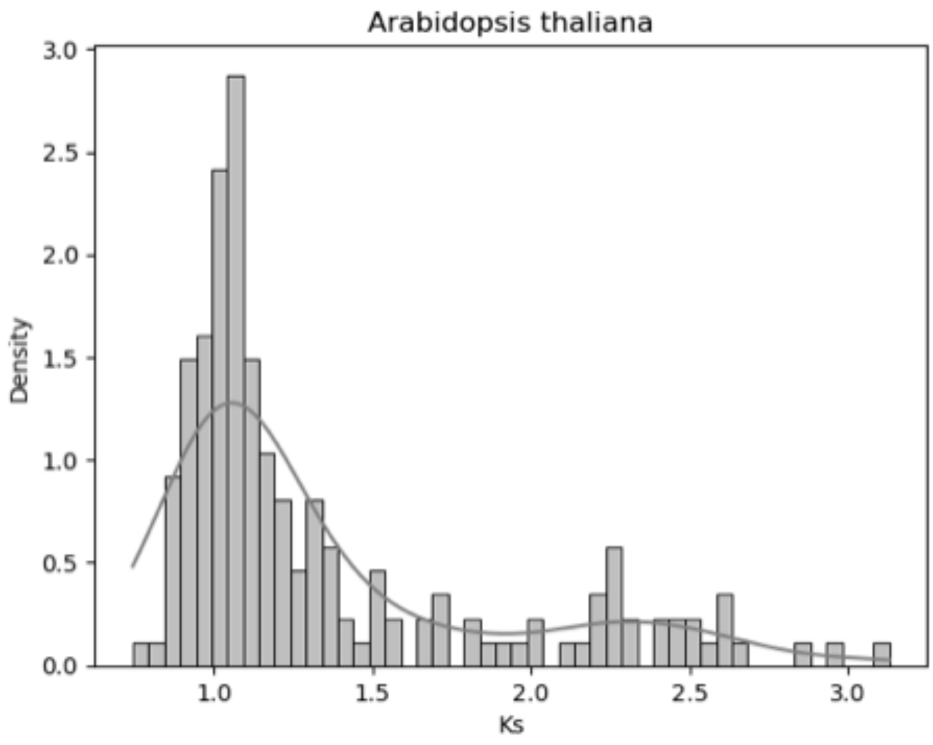


### Text S4. Snakefile_MCScanX_6species

#### species_mcscanx_list

`Purpose`: This step is to acquire a list of species name for running the MCScanX.

`scripts`:

```
mkdir -p {params.dir1}; \
ls -l {params.dir}|awk '{{print $9}}'| tee {params.dir1}/list.txt;
\
```

`Output`: "data/mcscanx/{ks_test_name}/list.txt"

```
>NP_001030613.1
MLLSALLTSVGINLGLCFLFFTLYSILRKQPSNVTVYGPRLVKKDGKSQQSNEFNLERLLPTAGWVKRA
LGLDALVFIRVFVFSIRVFSFASVVGIFILLPVNYMGTEFEEFFDLPKKSMDNFSISNVNDGSNKLWIH
```

#### species_mcscanx_file

`Purpose`: this is to move the respective input files into MCScanX folder for the Six species demonstrated in the protocol.

`scripts`:

```
while read line; do \
cp {params.dir}/$line/*.gff {params.dir1}||true; \
cp {params.dir}/$line/*.blast {params.dir1}||true; \
cp {params.dir}/$line/*.cds {params.dir1}||true; \
done < {input}; \
cat {params.dir1}/*.gff|tee {output.gff}; \
cat {params.dir1}/*.blast|tee {output.blast}; \
cat {params.dir1}/*.cds|tee {output.cds}; \
```

`Output`:

- gff = "data/mcscanx/{ks_test_name}/{ks_test_name}.gff",
- blast = "data/mcscanx/{ks_test_name}/{ks_test_name}.blast",
- cds = "data/mcscanx/{ks_test_name}/{ks_test_name}.cds"

#### The rest steps are similair to the previous file. But will take longer time ti run, since there are six species running together here.
